# Blockade of β-adrenergic receptor signaling improves cancer vaccine efficacy through its effect on naive CD8^+^ T-cell priming

**DOI:** 10.1101/497263

**Authors:** Clara Daher, Lene Vimeux, Ralitsa Stoeva, Elisa Peranzoni, Georges Bismuth, Emmanuel Donnadieu, Nadège Bercovici, Alain Trautmann, Vincent Feuillet

## Abstract

β-adrenergic receptor (β-AR) signaling, by acting directly on tumor cells and angiogenesis, has been showed to exert pro-tumoral effects. Growing evidence also suggests that β-AR expressed by immune cells affect the associated anti-tumor immune response. However, how and where β-AR signaling impinges the anti-tumor immune response is still unclear. Using a mouse model of vaccine-based immunotherapy, we show here that propranolol, a non-selective β-blocker, strongly improved the efficacy of the vaccine by enhancing the frequency of CD8^+^ T lymphocytes infiltrating the tumor (TILs). However, propranolol had no obvious effect on the reactivity of CD8^+^ TILs, a result further strengthened by *ex-vivo* experiments showing that these cells are insensitive to AR signaling triggered by adrenaline or noradrenaline. In contrast, we show that naive CD8^+^ T cell activation was strongly inhibited by β-AR signaling and that the beneficial effect of propranolol mainly occurred during their initial priming phase. We also demonstrate that the differential sensitivity of CD8^+^ TILs and naive CD8^+^ T cells is related to their activation status since *in* vitro-activated CD8^+^ T cells behaved similarly to CD8^+^ TILs, both exhibiting a down-regulation of the β_2_-AR expression. These results reveal that the initial priming phase of the anti-tumor response in the tumor-draining lymph node is a decisive part of the suppressive effect of β-AR signaling on the CD8^+^ T-cell response against cancer. These findings provide a rationale for the strategic use of clinically available β-blockers in patients to improve cancer immunotherapies such as anti-cancer vaccination strategies.

## Introduction

Multiple immunosuppressive mechanisms within the tumor microenvironment (TME) are known to alter the establishment of effective anti-tumor immune responses. They include cellular components such as regulatory CD4^+^ T cells (Tregs) (1), physical barriers imposed by the extracellular matrix (2), inflammatory mediators (*e.g.,* prostaglandin E2 (PGE2), adenosine) (3,4) and enhanced expression of inhibitory immune checkpoints (5). Therefore, current immune-based therapies, such as cancer vaccination, adoptive immunotherapy and immune checkpoint blockade, aim at shifting the balance towards more activated T cells and less suppressive mechanisms (6). Nevertheless, despite promising clinical results in various types of cancer, only a fraction of patients responds to immunotherapies, and many results indicate that combinations with treatments targeting immunosuppressive molecules, such as PGE2 and adenosine, within the TME could improve their efficiency (4,7).

Over the last decade, due to novel findings linking adrenergic receptor (AR) signaling and tumor progression (8), the nervous sympathetic system has emerged as an important player in cancer. The physiological impact of this system largely depends on the release of endogenous catecholamines, adrenaline and noradrenaline, from the adrenal medulla and the peripheral neurons innervating tissues under basal conditions, a secretion increased in response to a psychological stress. Retrospective epidemiological studies have shown that behavioral factors (*e.g.,* anxiety, depression) can influence the incidence and evolution of several types of cancers (9,10), and that patients treated with β-AR antagonists, referred as β-blockers, have a significantly better overall survival (11–13). Moreover, in mouse tumor models, studies have demonstrated that β-AR signaling favors tumor growth and metastatic invasion by acting on tumor cells, on angiogenesis and on the reshaping of the tumor stroma (14–16). Besides, growing evidence suggests that adrenergic signals can also exert their suppressive effects on anti-tumor immune responses by acting directly on immune cells that express functional ARs (17).

Although immune cells express both α- and β-ARs, it is generally admitted that β_2_-AR is the most highly expressed AR subtype, particularly in T and B cells. β_2_-ARs belong to the G protein-coupled receptor (GPCR) family and are coupled to Gα_s_ protein. In the canonical pathway, β_2_-AR signaling activates downstream transcription factors in a cAMP-Protein Kinase A (PKA)-dependent manner (18). Numerous reports have shown that β_2_-AR signaling modulates the functions of natural killer (NK) cells, innate lymphoid cells, dendritic cells (DCs), macrophages, B cells and T cells (18). Thus, in a cancer context, β-AR signaling has been shown to promote tumor growth or metastasis through M2 polarization of macrophages (19,20), infiltration of immunosuppressive cells such as Tregs and myeloid-derived suppressor cells (MDSCs) (21,22) and suppression of NK cell activity (23,24). Recent studies have also shown that β-AR signaling prevents the infiltration of CD8^+^ T cells in the tumor and could suppress their functions by acting either directly on CD8^+^ T cells (25–27) or indirectly on DCs (28) and MDSCs (21).

The objective of this study was to investigate how β-AR signaling could modify an anti-tumor CD8^+^ T cell-dependent response, and in particular whether this signaling mostly affects the priming or the effector phase of this response, or both. To address these questions, we used a murine model of vaccine-based tumor immunotherapy, in which CD8^+^ T cells were previously described to play a major role in tumor regression (29,30). We demonstrate here that propranolol, a non-selective β-blocker, strongly improved the efficacy of the vaccine by enhancing CD8^+^ T-cell tumor infiltration, but we did not observe any obvious effect of propranolol on the reactivity of tumor infiltrating CD8^+^ T cells (TILs). This was reinforced by *ex vivo* experiments showing that CD8^+^ TILs were insensitive to adrenaline/noradrenaline treatment. On the contrary, we show that purified naive CD8^+^ T cells were strongly sensitive to β-AR signaling, and that the beneficial effect of β–blockers mainly occurs during the initial priming phase of CD8^+^ T cells in the tumor-draining lymph node (TDLN). Finally, we show that the differential sensitivity of CD8^+^ TILs and naive CD8^+^ T cells is related to their activation status, since *in* vitro-activated CD8^+^ T cells behave similarly to CD8^+^ TILs, as they both exhibit a down-regulation of the β_2_-AR expression.

## Materials and Methods

### Mice

C57BL/6 and OT-1xSJL mice were purchased from Charles Rivers and maintained in the Cochin Institute Specific-Pathogen-Free animal facility. All experiments were performed in accordance with the Federation of European Laboratory Animal Science Associations. All procedures were approved by the Paris-Descartes Ethical Committee for Animal Experimentation (approval 03506.02).

### Chemicals

L-(−)Epinephrine-(+)-bitartrate (Adrenaline) and L-(−)Norepinephrine-(+)-bitartrate (Noradrenaline) were purchased from Calbiochem, Prostaglandin E2 (PGE2) from Tocris Bioscience and (±)-Propranolol hydrochloride from Sigma-Aldrich.

### Tumor model and *in vivo* treatments

The E7 expressing-TC1 (Tissue Culture 1) tumor cell line was maintained in complete RPMI (RMPI Medium 1640 GlutaMax, Gibco), including 10% fetal calf serum (FCS, GE Healthcare), Penicillin 50U/ml (Gibco), 50 μg/ml Streptomycin (Gibco) and 1 mM Sodium Pyruvate (Gibco).

Eight weeks-old C57BL/6J mice were subcutaneously injected on the flank with 1 × 10^5^ E7 expressing-TC1 tumor cells in 100 μL PBS. When tumors reached a volume of 100 mm^3^ (~10 days), mice were primed (day 0) with a peritumoral injection of 20 μg of STxBE7 vaccine (29), composed of the non-toxic β-subunit of Shiga toxin coupled to HPV16 derived-E7 peptide, and 6 × 10^5^ U of IFN-α in a total volume of 200 μL. The STxBE7 vaccine was provided by Dr Eric Tartour (Hôpital Européen Georges Pompidou, Paris, France) and Dr Ludger Johannes (Institut Curie, Paris, France). IFN-α was a kind gift from Dr Agnès Le Bon (Institut Cochin, Paris, France). The non-vaccinated mice were injected with PBS as control. The next day, vaccinated mice received the same dose of IFN-α. Mice were daily treated with propranolol (drinking water, 0.5 mg/mL) commencing one day prior to vaccination (d-1). Drinking water was changed two times per week. For some experiments, propranolol treatment was given to non-vaccinated mice 9 days after tumor implantation and to vaccinated mice 4 days after the vaccine prime. Tumors were measured every 3 days, and the tumor volumes were estimated by tumor volume = width × width × length/2. Tumors were harvested 10 days post-priming for flow cytometry, transcriptomic analysis and immunofluorescence.

### Tumor and lymph node cell suspensions

Fresh TC1 tumors were cut into 2-3 mm pieces and were digested for 45 min at 37°C with DNaseI (100 μg/ml, Roche) and collagenase (1 mg/ml, Roche) in RPMI medium. Red blood cells were lysed using ACK lysing buffer (Thermofisher Scientific) for 2-3 min at room temperature (RT). Cell suspensions of tumors were passed through a 40-μm nylon cell strainer (Corning), washed and resuspended in PBS, 2% FCS, 1 mM EDTA (Gibco). Lymph nodes (LNs) were processed into singlecell suspensions by mechanical disruption directly through a 40-μm nylon cell strainer.

### Flow cytometry

After washing in PBS, cell suspensions (2-3 × 10^6^) were stained in 96-wells round bottom plates with Live/Dead fixable dead cell stain kit (Invitrogen) for 20 min on ice, to gate out dead cells. Fc receptors were blocked with mouse Fc block (BD Biosciences) for 20 min on ice. Cells were then washed, resuspended in staining buffer (PBS, 2% FSC, 1 mM EDTA) and labeled for 20 min on ice with the following fluorescently conjugated anti-mouse antibodies (Abs): anti-CD45 (30-F11), anti-CD11b (M1/70), anti-CD11c (N418), anti-Ly6G (1A8), anti-Ly6C (HK1.4), anti-F4/80 (BM8), anti-CD64 (X54-5/7.1), anti-XCR1 (REA707), anti-CD172a (P84), anti-MHC class II (M5/114.15.2), anti-TCRβ (H57-597), anti-TCRγδ (GL3), anti-CD4 (GK1.5), anti-CD8a (53-6.7), anti-CD19 (1D3), anti-NK1.1 (PK136), anti-CD69 (H1.2F3), anti-PD-1 (29F.1A12), anti-CD44 (IM7), and anti-CD62L (MEL-14). For detection of E7-specific CD8^+^ T cells, cells were stained with Kb-E7 dextramer (RAHYNIVTF peptide; Immudex) for 30 min on ice. After staining, cells were washed in staining buffer and were fixed in 2% PFA (Electron Microscopy Sciences) for 15 min at RT. For intracellular staining, cells were fixed and permeabilized using Foxp3/Transcription staining buffer set (eBioscience) or Cytofix/Cytoperm, solution (BD Biosciences) as per the manufacturer’s instructions. Cells were labeled overnight at 4°C with the following fluorescently conjugated anti-mouse Abs: anti-Foxp3 (FJK16S), anti-ki-67 (polyclonal), anti-granzyme B (GzmB; GB11) and anti-IFN-γ (XGM1.2). For intracellular IFN-γ staining, cells were stimulated *in vitro* for 4 h at 37°C with either 5 μg/ml E7-peptide (kindly provided by Dr Eric Tartour) or Dynabeads Mouse T-activator CD3/CD28 (1 bead for 2 CD8^+^ T cells; Thermofisher) or with 0. 5 μg/mL PMA + 2.5 μM Ionomycin (Calbiochem) in the presence of Brefeldin A (GolgiPlug, BD Bioscience). Fluorescent Abs were purchased either from BD Pharmingen, BioLegend, eBioscience, Miltenyi Biotech or EMD Millipore. Cells were acquired on LSR II flow cytometer (BD Biosciences) or Fortessa flow cytometer (BD Biosciences) and analyzed using FlowJo v10 software (FlowJo LLC).

### CD8^+^ T-cell isolation and culture

CD8^+^ T cells and CD8^+^ OT-I cells were purified from spleen and LNs from C57BL/6J mice using a negative isolation kit (Invitrogen) as per as the manufacturer’s instructions. The CD8^+^ T cell enriched fraction was labeled with anti-CD8a, anti-CD44 and CD62L fluorescent Abs and naive CD8^+^ T cells (CD8^+^CD44^nes^CD62L^+^) were sorted on ARIA III (BD Biosciences). Activated CD8^+^ T cells were obtained from CD8^+^ T cells activated for 7 days with plate-bound anti-CD3/CD28 Abs (2 μg/mL each; 145-2C11/37.51; BD Biosciences), with a 1:2 split on days 3 and 5 in complete RPMI supplemented with IL-2 (20 U/ml).

For CD8^+^ TILs isolation, cell suspensions from vaccinated TC1 tumor-bearing mice were resuspended in 40% Percoll (GE Healthcare) layered on top of 80% Percoll, and centrifuged at 325 g for 25 min without brake. Cells at the Percoll interface were collected, labeled with anti-CD45, anti-CD4, anti-CD8a, anti-CD11 b fluorescent Abs and CD8^+^ TILs (CD45^+^CD11b^nes^CD4^nes^CD8^+^) were sorted on FACSARIA III (BD Biosciences). CD8^+^ TILs were either used immediately (RNA extraction) or let to recover 24 h in complete RPMI supplemented with IL-2 (20 U/ml).

### Calcium influx measurements

T cells were purified from LNs using an EasySep T-cell isolation kit (Stemcell technologies) as per as the manufacturer’s instructions. Cells were loaded for 30 min at 37°C in complete RPMI with the membrane-permeable fluorescent Ca^2+^ indicator dye Indo-1 AM (5 μM; Invitrogen), washed and labeled with anti-CD4, anti-CD8a, anti-CD62L and anti-CD44 fluorescent Abs for 20 min on ice. After washing in complete RPMI, T cells (2 × 10^6^) were rested for 5 min and then pre-treated with either adrenaline, noradrenaline or PGE_2_ for 10 min at 37°C. In some experiments, T cells were pre-treated with propranolol for 30 min at 37°C prior to adrenaline or noradrenaline treatment. Changes in intracellular Ca^2+^ (iCa^2+^) concentration over the time were monitored with the Indo-1 iCa^2+^-free (violet) and Indo-1 iCa^2+^-bound (blue) channels on a LSR II flow cytometer at 37°C. The calcium baseline level was measured for 30 s before stimulation with hamster anti-CD3ε Abs (2.5 μg/ml; 145-2C11; BD Biosciences) and then for 1 min before cross-linking with goat anti-hamster IgG secondary Abs (20 μg/ml; Invitrogen). Peaks of iCa^2+^ mobilization were quantified in naive CD8^+^ T cells (CD8^+^CD44^nes^CD62L^+^) as the ratio of Indo-1 iCa^2+^-free /Indo-1 iCa^2+^-bound. Calcium responses were also assessed in Facs-sorted CD8^+^ TILs (2 × 10^5^) and in activated CD8^+^ T cells (2 × 10^6^) following similar protocol.

### Proliferation and survival assay

Naive CD8^+^ T cells were labeled with 5 μM Celltrace Violet (CTV; Invitrogen) according to manufacturer’s instructions and stimulated with plate-bound anti-CD3/CD28 Abs. Cells (5 × 10^4^) were cultured with or without IL-2 (20 U/ml) in the presence of adrenaline or noradrenaline. Catecholamine treatments were renewed at 24 h and proliferation was assessed by flow cytometry after 3 days. Proliferation index represents the total number of divisions divided by the number of cells that went into division and was calculated using the following formula: proliferation index 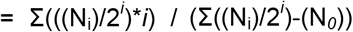 where N is the cell number and *i* is the generation number (parent generation = 0). Facs-sorted CD8^+^ TILs or activated CD8^+^ T cells (5 × 10^4^) were cultured with or without IL-2 (20 U/ml) in the presence of adrenaline or noradrenaline during 24 h. Cell populations were enumerated using Accuri C6 flow cytometer (BD Biosciences) at days 0 and 1. Proliferation rates were quantified as the ratio of absolute cell number at D1/ absolute cell number at D0. To assess effects of catecholamines on CD8^+^ T cells viability, naive CD8^+^ T cells (5 × 10^4^) were cultured with IL-7 (10 ng/ml, PeproTech) and Facs-sorted CD8^+^ TILs (5 × 10^4^) were cultured with or without IL-2 (20 U/ml). Survival was evaluated with Live/Dead staining by flow cytometry.

### Generation of bone marrow–derived DCs (BMDCs)

Bone marrow cells were collected from femurs of C57BL/6J mice, depleted of red blood cells using ACK buffer, and were cultured at 1 × 10^7^ cells per well in tissue-culture-treated 6-well plates in 4 ml of complete RPMI and 20 ng/ml GM-CSF (BioLegend). Half of the medium was removed at day 2 and new medium warmed at 37°C and supplemented with 40 ng/ml GM-CSF was added. The culture medium was entirely discarded at day 4 and replaced by fresh warmed medium with 20 ng/ml GM-CSF. On day 6, non-adherent cells in the culture supernatant and loosely adherent cells harvested by gentle washing with PBS were pooled and used for experiments.

### BMDC maturation and coculture with T cells

Immature BMDCs were stimulated with 2.5 μg/mL of STxBOVA vaccine and 10^5^ U/ml of IFN-α in the presence of adrenaline or noradrenaline for 18 h. The STxBOVA vaccine was provided by Dr Eric Tartour (Hôpital Européen Georges Pompidou, Paris, France) and Dr Ludger Johannes (Institut Curie, Paris, France). For DC maturation analysis, BMDCs were collected and Fc receptors were blocked with mouse Fc block for 20 min on ice. Cells were then stained with fluorochrome-conjugated Abs against surface markers CD11b, CD11c, MHC class II, CD80 and CD86 in staining buffer for 20 min on ice. For DC-OT-I coculture, 1 × 10^4^ stimulated BMDCs were cultured with 5 × 10^4^ purified OT-I CD8^+^ T cells labeled with 5 μM CTV. T cell proliferation was assessed by flow cytometry after 3 days.

### ELISA

IL-2 and IFN-γ cytokine production was measured in culture supernatant by ELISA Max kit (BioLegend) according to manufacturer’s indications.

### RNA extraction and real-time qRT-PCR

Total RNA from whole tumor or CD8^+^ T cells were extracted using RNeasy Mini Kit (Qiagen) according to the manufacturer’s instructions and gene expression was analyzed by RT-qPCR with the LighCycler 480 Real-Time PCR system (Roche Life Science). For cDNA synthesis, 0.2 to 1 μg of purified RNA was reverse-transcribed using the Advantage RT-for-PCR kit (Applied Clontech). PCR amplification reactions were carried out in a total volume of 10 μl using LighCycler 480 SYBR Green I Master (Roche Life Science), forward and reverse primers (10 μM each) (sequences are detailed in **Supplementary Table S1**) and cDNA. GAPDH was used as a housekeeping gene to normalize mRNA expression. Relative expression was calculated by 2^-ΔΔCt^ method to compare expression levels of the same transcript in different samples or 2^-ΔCt^ to compare expression levels of several transcripts in the same sample. All the measures were performed in triplicate and validated when the difference in threshold cycle (Ct) between two measures was <0.3. Raw Ct values were calculated using the LightCycler 480 v1.5.0 SP3 software.

### Immunofluorescence

TDLNs of vaccinated non-treated TC1 tumor-bearing mice were fixed for 2 h at RT in a Periodate–Lysine–Paraformaldehyde solution (0.05 M phosphate buffer containing 0.1 M L-lysine (pH 7.4), 2 mg/mL NaIO4, and 10 mg/mL paraformaldehyde). Tissue slices were prepared as previously described (2). Briefly, fixed TDLNs were embedded in 5% low-gelling-temperature agarose (type VII-A; Sigma-Aldrich) prepared in PBS. Ganglionic slices (250 μm) were cut with a vibratome (VT 1000S; Leica) in a bath of ice-cold PBS. Immunostaining was performed by first blocking Fc receptors with mouse Fc block for 1 h at RT. After washing in PBS 0.3% Triton X-100 1% BSA 5% donkey serum, slices were stained for sympathetic nerves with anti-tyrosine hydroxylase primary Abs (rabbit polyclonal; EMD Millipore) overnight at 4°C. Slices were then washed and stained with PE-conjugated anti-CD3 Abs (eBioscience) for 30 min at RT. After washing, immunodetection was performed using Alexa fluor 647-conjugated donkey anti-rabbit IgG (H+L) secondary Abs (Life Technologies) for 30 min at RT. All Abs were diluted in PBS 0.3% Triton X-100 1% BSA 5% donkey serum. Images were obtained with a spinning disk confocal microscope (Leica) equipped with a CoolSnap HQ2 camera and a 25X water immersion objective (25X/0.95 N.A.). All images were acquired with MetaMorph 7 imaging software (Molecular Devices) and analyzed with ImageJ software.

### Statistical analysis

Data were analyzed with GraphPad Prism6 Software. For comparison of two non-parametric datasets, Mann-Whitney U-test was used. For multigroup comparisons, one-way ANOVA or two-way ANOVA tests with post hoc testing using Tukey’s multiple comparison were used. *P* values <0.05 were considered significant (*, *P* <0.05, **, *P* <0.01, ***, *P* <0.001, ****, *P* <0.0001). All data are depicted as Mean ± SEM.

## Results

### Blocking β-AR signaling improves vaccine-induced tumor regression

In order to elucidate the effects of the β-AR signaling on a CD8^+^ T cell-dependent anti-tumor immune response, we assessed the effects of the pan-β-AR antagonist propranolol on mice engrafted with TC1 tumor cells, and vaccinated with the STxBE7-vaccine plus IFN-α, to stimulate innate immunity. STxBE7-vaccine was previously described to induce the regression of the E7 expressing TC1 tumors (29,30). C57BL/6J mice were transplanted with TC1 tumor cells and, when tumor nodules reached 100 mm_3_ at day 10, treated with a peri-tumoral injection of STxBE7-vaccine (30). Mice were treated with propranolol daily, starting one day prior to STxBE7-vaccination (**Fig. 1A**). We designed a sub-optimal vaccination protocol in which a minority of tumors completely regressed (≈30%) and the others either progressed (≈10%), stabilized (≈10%) or partially regressed (≈50%) (**Fig. 1B, 1C**). Indeed, such a variable outcome is appropriate for evaluating if the propranolol treatment can modulate the efficiency of the induced anti-tumor response. As shown in **Fig. 1B**, propranolol strongly improved the efficiency of STxBE7-vaccine. All the tumors from the propranolol-treated mice regressed, with 74% of the mice being tumor-free at the sacrifice 22 days after the vaccination and the others at least partially regressing (**Fig. 1C**). Notably, treatment of unvaccinated control mice with propranolol had no effect on tumor growth in this tumor model (**Fig. 1B**), suggesting that propranolol acts mainly on the antitumor immune response induced by the vaccine. These results demonstrate that blocking β-AR signaling strongly improves the efficacy of an anti-tumor response induced by a vaccine-based immunotherapy.

**Figure 1.**
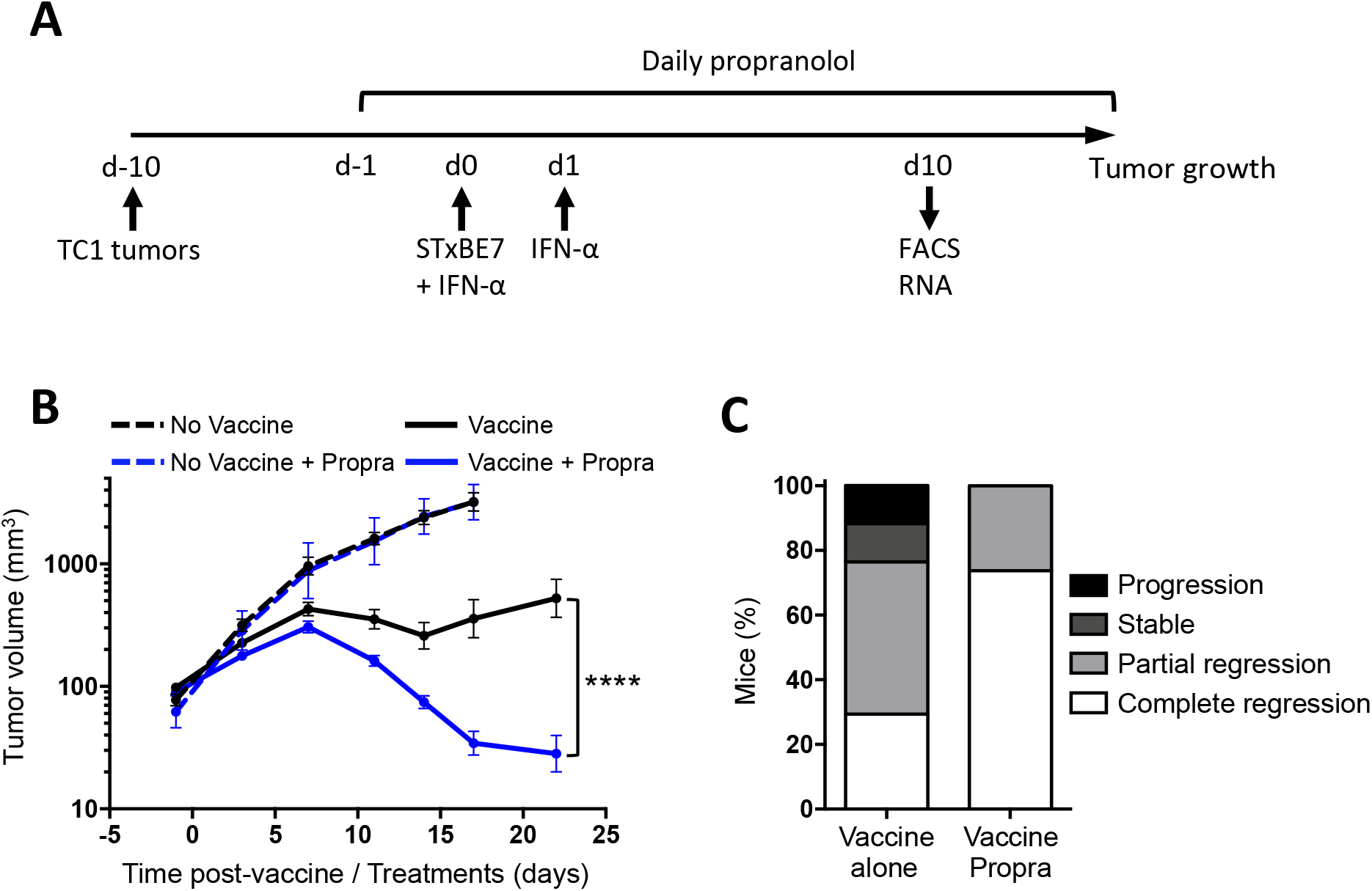
Propranolol treatment improves vaccine-induced tumor regression. C57BL/6J mice were s.c. injected with 1 × 10^5^ E7 expressing-TCl tumor cells. STxBE7 vaccine plus IFN-α were administrated ten days later (d0). The next day, vaccinated mice received an additional dose of IFN-α. Mice were daily treated with propranolol commencing one day prior to vaccination (d-1). **A**, Experimental design. **B**, Tumor growth curves of non-vaccinated non-treated mice (dashed black line), non-vaccinated propranolol-treated mice (dashed blue line), vaccinated non-treated mice (black line) and vaccinated propranolol-treated mice (blue line). **C**, Repartition of vaccinated non-treated mice and of vaccinated propranolol-treated mice. Results were grouped as progression (no regression), stable (tumor size decreased by < 30% from day 7), partial regression (tumor size decreased by > 30% from day 7) or complete regression (tumor free at sacrifice day 22). Pool from three independent experiments (*n* = 17 to 19 per group). Statistical analysis by two-way ANOVA test: ****, *p* <0.0001. Mean ± SEM.

### Blocking β-AR signaling increases the frequency of CD8^+^ T cells infiltrating the tumor without affecting their reactivity

To investigate why we observed a better tumor regression after propranolol treatment and which immune cells of the TME are affected by the treatment, immune intra-tumoral infiltrates were analyzed 10 days post-vaccination by multi-parametric flow cytometry. The gating strategies of the different lymphoid and myeloid cell populations are shown in **Supplementary Fig. S1A and S1B**. As expected, vaccination led to a strong infiltration of CD45+ immune cells into the tumor (**Fig. 2A**). This concerned all lymphoid and myeloid cell subsets, except conventional DCs (cDCs), the frequency of which was decreased by vaccination (**Supplementary Fig. S2A**). Moreover, a careful look at the repartition of the different immune populations among CD45+ immune cells showed that the vaccination induces a profile of anti-tumor response, characterized by an increased frequency of CD8^+^ T cells, NK cells, monocytes and a decrease in Tregs and tumor-associated macrophages (TAMs) (**Supplementary Fig. S2B**). In vaccinated mice, propranolol significantly increased the abundance of CD45+ immune cells (**Fig. 2A**). Most importantly, only CD8^+^ T cells, including the E7-specific ones, were significantly enriched (≈ 2.5-fold) in the tumor and among CD45+ immune cells by propranolol treatment (**Fig. 2B and Supplementary Fig. S2A, S2B**).

**Figure 2.**
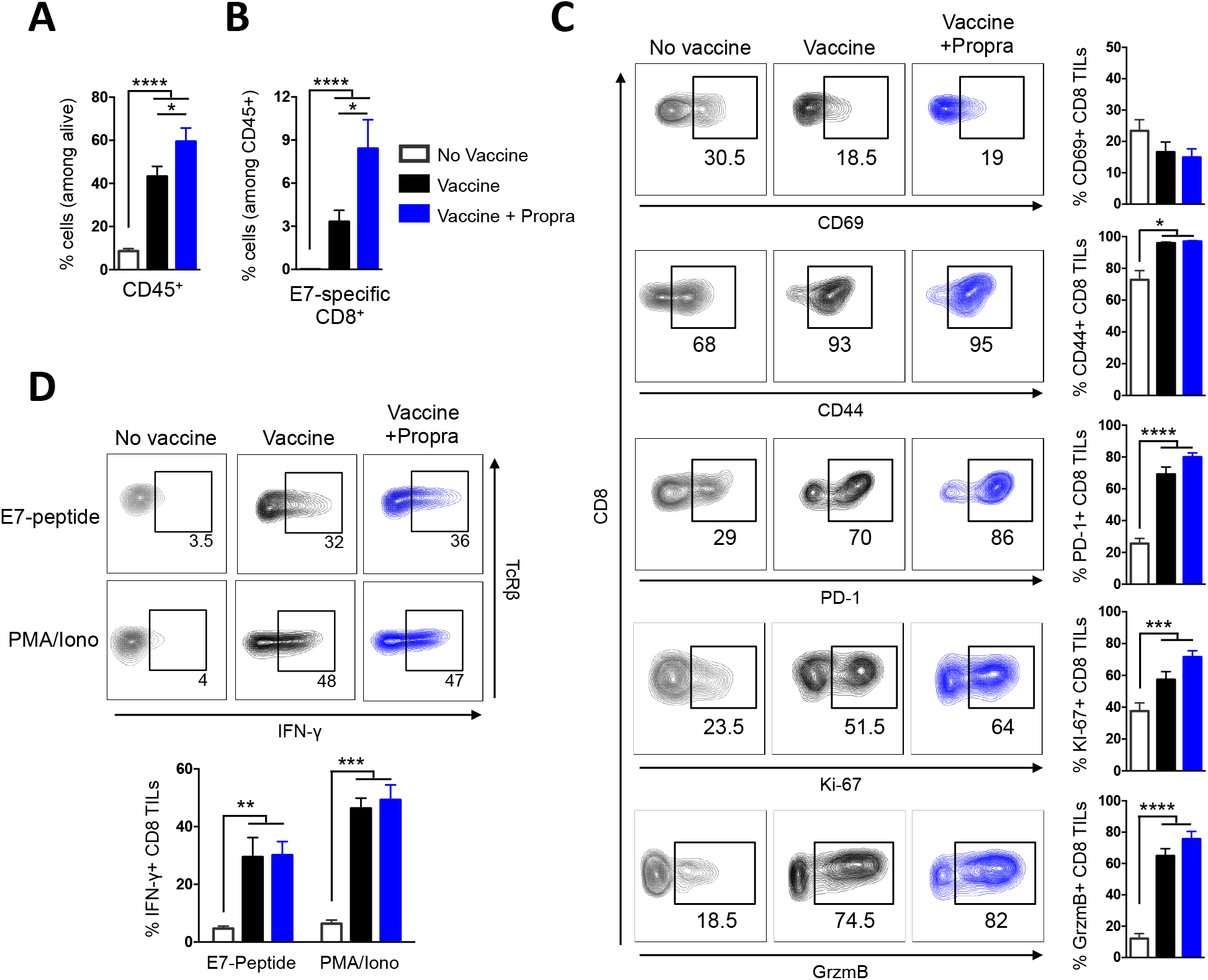
Propranolol treatment increases the frequency of CD8^+^ TILs without affecting their reactivity. C57BL/6J mice were s.c. injected with E7 expressing-TC1 tumor cells and vaccinated as mentioned in Figure 1. Day 10 post-vaccination, immune cell infiltrates from tumor single-cell suspensions were analyzed by flow cytometry. **A**, Frequency of CD45^+^-cells among live cells. **B**, Frequency of E7-specific CD8^+^ T cells among CD45^+^-cells. **A** and **B**, Pool from six independent experiments (*n* = 16 to 18 per group). **C**, Expression of activation markers CD69, CD44, PD-1, Ki-67 and GrzmB on CD8^+^ TILs, with representative flow plots of markers expression in CD8^+^ TILs. Pool from four independent experiments (*n* = 12 to 15 per group). **D**, Percentage of IFN-γ^+^ CD8^+^ TILs following *in vitro* restimulation with 5 μg/ml E7-peptide or PMA/ionomycin for 4 h, with representative flow plots of IFN-γ staining in CD8^+^ TILs. Pool from two independent experiments (*n* = 8 to 12 per group). Statistical analysis by Mann-Whitney test: *, *P* <0.05, **, *P* <0.01, ***, *P* <0.001, ****, *P* <0.0001. Mean ± SEM.

We next wondered whether the reactivity of CD8^+^ TILs was improved by β-AR signaling blockade, and thus analyzed the expression of the activation markers CD69, CD44 and PD-1, and the percentage of cells producing Granzyme B or positive for the proliferation marker Ki-67. Except for CD69 expression, vaccination led to an increase of all these activation markers (**Fig. 2C**). Most importantly, although we found an increased frequency of CD8^+^ TILs in the propranolol-treated mice, their activation status was comparable to that of CD8^+^ TILs in untreated vaccinated mice (**Fig. 2C**). This phenotypic observation was reinforced by a functional one. Thus, we tested a potential propranolol influence on the ability of CD8^+^ TILs to produce IFN-γ after restimulation with either the E7-peptide or a PMA/ionomycin combination. For both conditions, vaccination clearly induced IFN-γ-producing CD8^+^ TILs, but there was no difference in the production of IFN-γ whether the vaccinated mice had been treated with propranolol or not (**Fig. 2D**).

These results suggest that propranolol treatment improves the efficacy of an anti-tumor vaccination by increasing the frequency of the CD8^+^ TILs without affecting their activation status or their ability to produce IFN-γ.

### Propranolol treatment improves CD8^+^ T cell priming in the tumor-draining lymph node

We next decided to gain insight into the potential mechanism underlying the increase in CD8^+^ TIL frequency induced by propranolol treatment. We first examined the transcripts of a series of chemokines in the whole tumor. In all cases, the major effect was an increase of their expression after vaccination. However, propranolol treatment had no significant effect on the transcripts of chemokines, in particular CXCL9 and CXCL10, which are major T cell attractants in such settings (31) (**Supplementary Fig. S3**). This shows that the effect of propranolol on the enhanced CD8^+^ T-cell infiltration within the tumor is not primarily chemokine-dependent.

Another possibility was that the increase of CD8^+^ TILs observed with propranolol treatment would be a consequence of its effect during their initial priming by antigen in the TDLN. This hypothesis was strengthened by the observation that the frequency of E7-specific CD8^+^ T cells in the TDLN, mainly controlled by the efficacy of tumor-specific T cell priming, was increased in propranolol-treated mice (**Fig. 3A**). Moreover, TDLN exhibits a sympathetic innervation (positive for tyrosine hydroxylase) in close contact with T cells (**Fig. 3B**), providing an anatomical evidence for a possible effect of β-AR signaling on the activation of naive CD8^+^ T cells. To test more directly this hypothesis, we designed a vaccination protocol in which β-AR blockade by propranolol was absent during the first 4 days following the vaccination, corresponding to the initial priming phase (**Fig. 3C**). In these conditions, we completely lost the beneficial effect of propranolol on the anti-tumor immune response, since vaccinated mice, treated or not with propranolol, showed very similar profiles of tumor regression (**Fig. 3D**). Consistently, such a delayed propranolol treatment also failed to increase the frequency of CD45+ cells and CD8^+^ T cells in the tumor (**Fig. 3E**).

**Figure 3.**
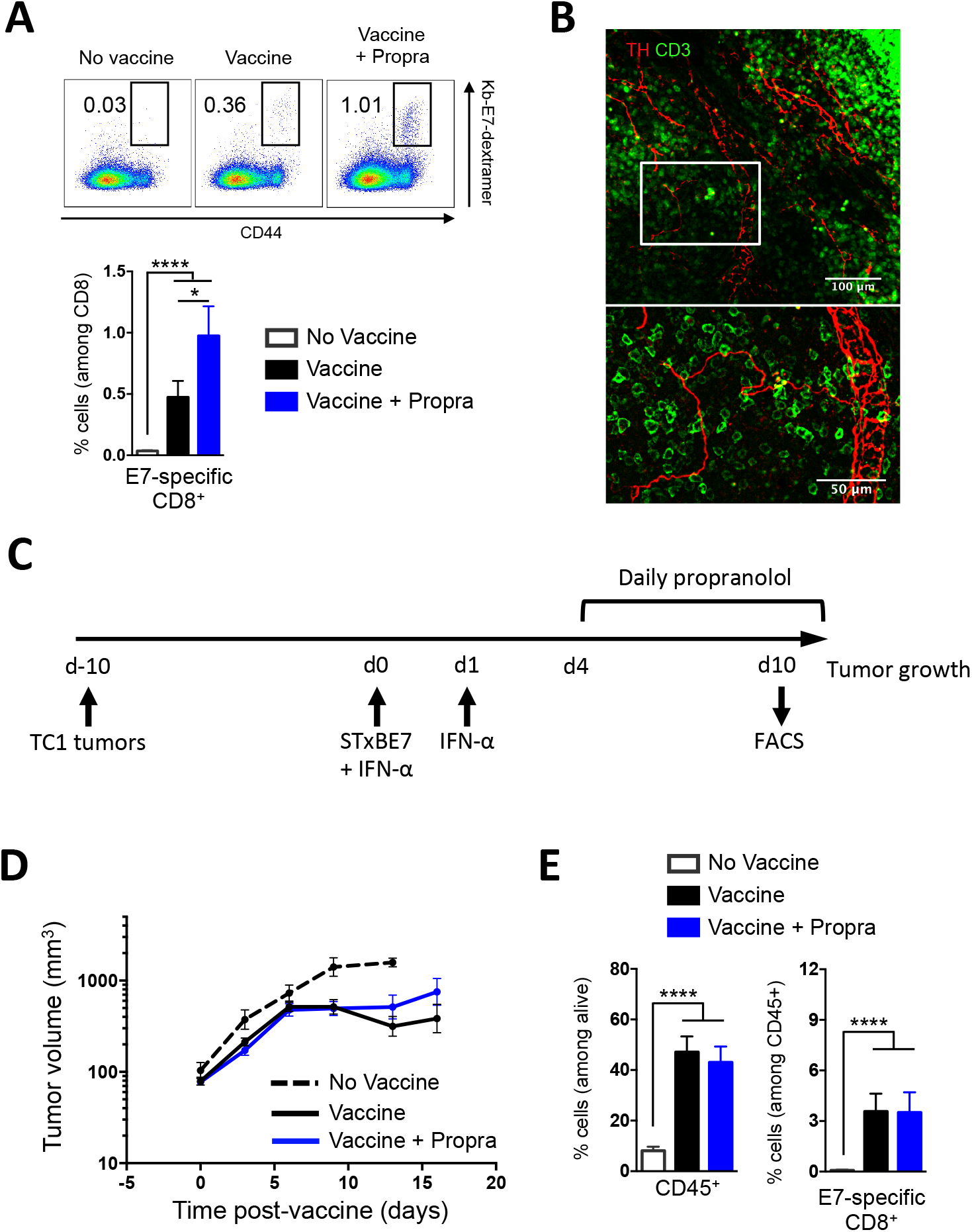
Propranolol treatment improves CD8^+^ T-cell priming in the TDLN. **A**, C57BL/6J mice were s.c. injected with E7 expressing-TCl tumor cells and vaccinated as mentioned in Figure 1. Frequency of E7-specific CD8^+^T cells from TDLN single-cell suspensions analyzed by flow cytometry at day 10 post-vaccination, with representative flow plots of Kb-E7-dextramer and CD44 staining in CD8^+^ T cells. Pool from three independent experiments (*n =* 15). **B**, Confocal image of a TDLN slice stained for tyrosine hydroxylase (TH, red) and CD3 (green). 25x magnification. C-E, C57BL/6J mice were s.c. injected with 1 × 10^5^ E7 expressing-TCl tumor cells. STxBE7 vaccine plus IFN-α were administrated ten days later (d0). The next day, vaccinated mice received an additional dose of IFN-α. Mice were daily treated with propranolol commencing 4 days after vaccination (d4). C, Experimental design. **D**, Tumor growth curves of non-vaccinated non-treated mice (dashed black line), vaccinated non-treated mice (black line) and vaccinated propranolol-treated mice (blue line). Pool from three independent experiments (*n* = 15). E, Frequencies of CD45^+^-cells among live cells and E7-specific CD8^+^ T cells among CD45^+^-cells from tumor single-cell suspensions analyzed by flow cytometry at day 10 post-vaccination. Pool from two independent experiments (*n =* 10). Statistical analysis by Mann-Whitney test: *, *P* <0.05, ****, *P* <0.0001. Mean ± SEM.

Altogether, these results demonstrate that the main effect of propranolol on the vaccine-induced tumor regression occurs during the priming phase in the TDLN by increasing the generation of tumor antigen-specific CD8^+^ T cells.

### AR signaling does not impair maturation and antigen presentation capacities of DCs stimulated with the vaccine

The aforementioned findings suggest that blockade of β-AR signaling improves the priming of the naive CD8^+^ T cells in the TDLN. To explain this result, a suppressive effect of AR signaling on the maturation and antigenic presentation capacity of DCs could be involved. To investigate this possibility, BMDCs were stimulated 18 h with an equivalent of the STxBE7-vaccine fused with OVA instead of E7 (STxBOVA-vaccine) plus IFN-α with or without the natural AR agonists, adrenaline and noradrenaline. Stimulation of DCs with STxBOVA-vaccine induced their maturation, as reflected by an increased expression of MHC class II and coreceptor molecules CD80 and CD86, which were not significantly reduced by agonist treatments (**Fig. 4A**). To test whether antigen presentation capacities of DCs were affected by adrenergic signals, BMDCs were pulsed with STxBOVA-vaccine plus IFN-α in the presence or absence of adrenaline and noradrenaline for 18 h, and then cocultured with OVA-specific CD8^+^ OT-I T cells. The DC-induced proliferation of OT-I was not affected by the treatment of DCs with AR agonists (**Fig. 4B**). Thus, adrenergic signaling does not impair maturation and antigenic presentation capacities of DCs stimulated with the vaccine.

**Figure 4.**
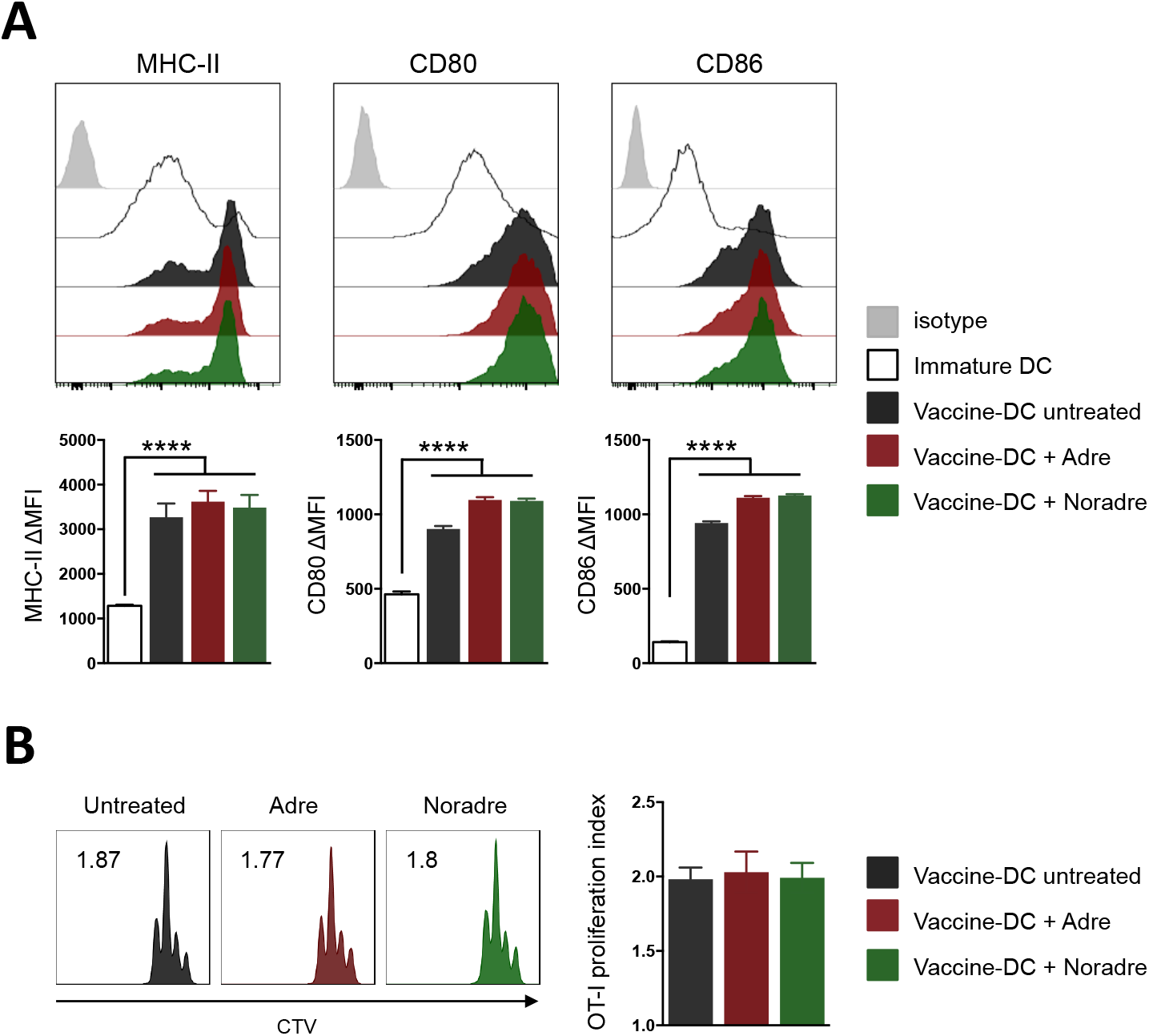
AR signaling did not alter vaccine-stimulated DC maturation and their antigen presentation capacity to naive CD8^+^ T cells. Immature BMDCs were pre-treated with either 10 μM adrenaline or noradrenaline for 1 h and stimulated with STxBOVA vaccine plus IFN-α for 18 h. A, Surface expression of MHC-II, CD80 and CD86 displayed as ΔMFI relative to isotype control (*n =* 8), with representative histograms of staining of BMDCs. B, Proliferation of CVT-labeled CD8^+^ OT-I cells co-cultured with pre-treated DCs for 3 days, expressed as proliferation index (*n* = 6), with representative histograms of CTV dilution in CD8^+^ OT-I cells. For all panel, one representative experiment of two is shown. Statistical analysis by one-way ANOVA test: ****, *p* <0.0001. Mean ± SEM.

### Naive CD8^+^ T cells are strongly susceptible to β-AR signaling

We next wondered if blockade of β-AR signaling could directly act on naive CD8^+^ T cells to improve their priming. To test this hypothesis, we evaluated the effect of treatments with adrenaline or noradrenaline on naive CD8^+^ T cell functions. The early calcium signaling response following CD3 stimulation was first analyzed. As shown in **Fig. 5A**, both agonists inhibited the anti-CD3-induced calcium response of naive CD8^+^ T cells in a dose-dependent manner. Adrenaline had the strongest effect, which probably reflects its higher affinity for β_2_-ARs, highly expressed in naive CD8^+^ T cells (**Supplementary Fig S4A**). The key role of β-AR signaling in this effect was demonstrated by the reversion of the inhibition by propranolol pre-treatment (**Fig. 5B**). We next assessed the consequences of this inhibitory effect in terms of T cell proliferation. As shown in **Fig. 5C**, the proliferation of naive CD8^+^ T cells at day 3 was reduced by agonist treatment in a dose-dependent manner. The inhibition was not due to an effect on naive CD8^+^ T cells viability, which was not affected by agonist treatments (**Supplementary Fig S4B**). However, proliferation was rescued by addition of IL-2 in the culture medium (**Fig. 5D**), suggesting that the inhibition was a consequence of reduced IL-2 production, as further confirmed by measuring IL-2 levels in supernatants after agonist treatment (**Fig. 5E**). In addition, we observed that adrenaline and noradrenaline suppressed IFN-γ secretion, measured in the supernatant 24 h after stimulation, i.e., before the cells had started to proliferate (**Fig. 5F**). Altogether, these results demonstrate that activation of naive CD8^+^ T cells is strongly inhibited by β-AR signaling, a phenomenon that may affect their priming in the TDLN.

**Figure 5.**
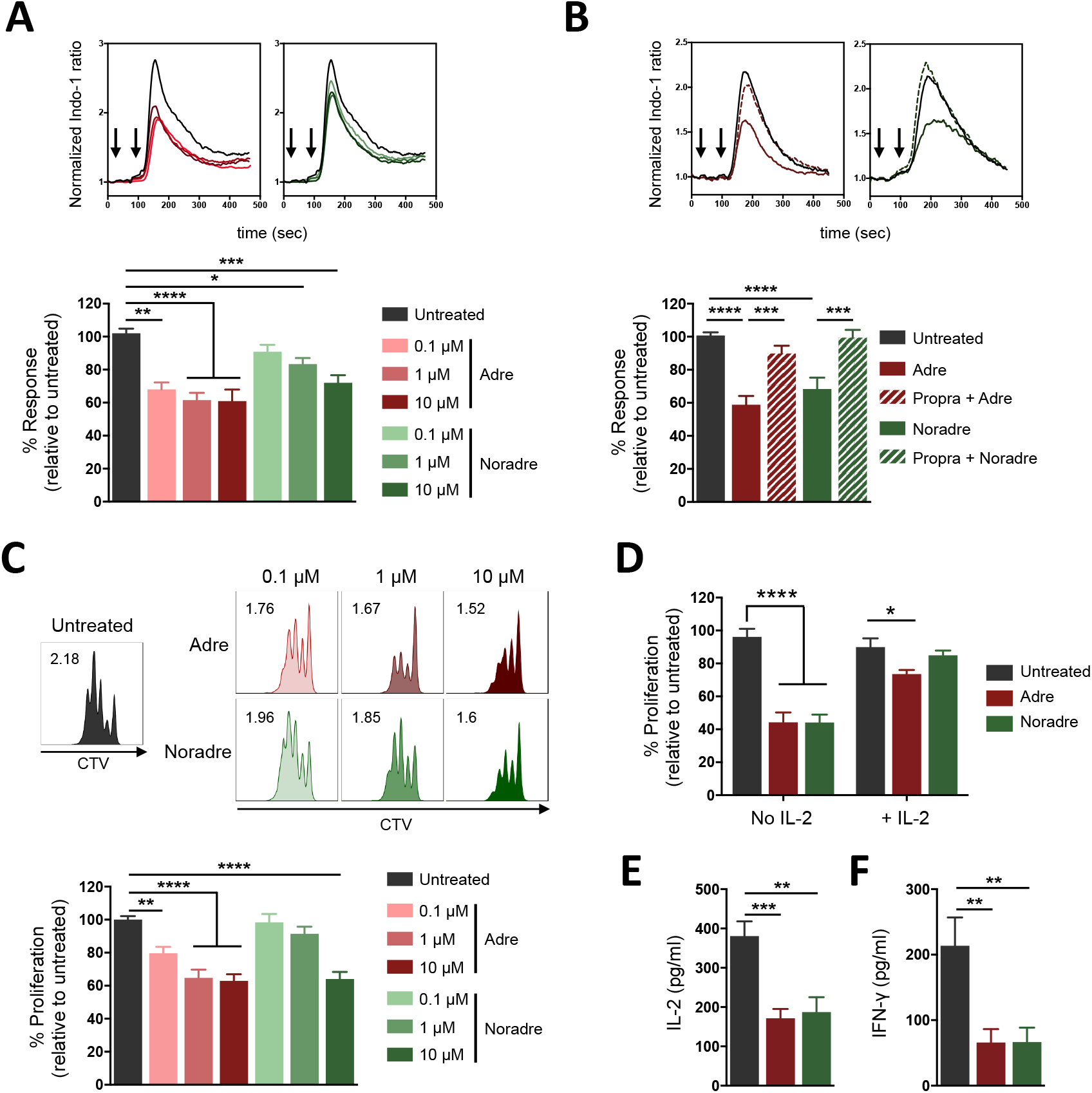
Naive CD8^+^ T cells are susceptible to β-AR signaling. **A** and **B**, Naive CD8^+^ T cells were loaded with Indo-1, pre-treated with either adrenaline or noradrenaline at the indicated concentration during 10 min (A) or pre-treated with 10 μM propranolol during 30 min followed by treatment with either 10 μM adrenaline or noradrenaline during 10 min (B). iCa^2+^ mobilization was assessed by flow cytometry before and after stimulation with hamster anti-CD3ε Abs (first arrow, above) cross-linked with antihamster IgG (second arrow) and is indicated as % response in treated cells relative to untreated cells, with representative normalized Indo-1 ratio histograms. **A**, Pool from four independent experiments (*n* = 4). **B**, Pool from two independent experiments (*n =* 5). C and D, Proliferation index of CVT-labeled naive CD8^+^ T cells stimulated with anti-CD3/CD28 Abs in the presence of adrenaline or noradrenaline at the indicated doses (C) or in the presence of 10 μM adrenaline or noradrenaline with or without IL-2 **(D)** during 3 days, indicated as % proliferation of treated cells relative to untreated cells, with representative histograms of CTV dilution. **C**, Pool from four independent experiments (*n =* 20). **D**, Pool from two independent experiments (*n =* 8). **E** and **F**, IL-2 **(E)** and IFN-γ **(F)** measured in the supernatants 24 h after stimulation of naive CD8^+^ T cells with anti-CD3/CD28 Abs in the presence of 10 μM adrenaline or noradrenaline. Pool from six independent experiments (*n* = 8). Statistical analysis by one-way ANOVA test: *, *P* <0.05, **, *P* <0.01, ***, *P* <0.001, ****, *p* <0.0001. Mean ± SEM.

### CD8^+^ TILs are not affected by adrenergic signals

Given the strong effects of β-AR signaling on naive CD8^+^ T-cell activation, we were surprised not to observe any obvious effect of propranolol treatment on CD8^+^ TIL activation and on their ability to secrete IFN-γ (**Fig. 2B, 2C**). We thus investigated how Facs-sorted E7-specific CD8^+^ TILs could react to adrenaline and noradrenaline *ex vivo.* These cells exhibited an activated/effector phenotype (CD44+CD62L^neg^) (**Supplementary Fig. S5A**). Surprisingly, contrary to what was observed with naive CD8^+^ T cells, their anti-CD3-induced calcium response was not affected by adrenaline and noradrenaline treatment (**Fig. 6A**). In contrast, PGE2, another cAMP-elevating GPCR ligand, previously described to inhibit T cell functions (32), strongly inhibited the response, eliminating the possibility of an unresponsiveness of these cells to the inhibitory effect of cAMP (**Fig. 6A**). We also found no effect of adrenaline and noradrenaline on the survival and proliferation of CD8^+^ TILs cultured with or without IL-2 (**Fig. 6B and Supplementary Fig. S5B**). This was consistent with the fact that we did not observe any effect of AR agonists on IL-2 production by CD8^+^ TILs restimulated with anti-CD3/anti-CD28 Abs (**Fig. 6C**). Finally, we showed that the production of IFN-γ by CD8^+^ TILs restimulated with anti-CD3/CD28 Abs was not inhibited by AR agonists (**Fig. 6D**). This was confirmed by IFN-γ intracellular staining experiments following restimulation of CD8^+^ TILs with either anti-CD3/CD28 Abs or PMA/ionomycin in the presence of adrenaline or noradrenaline during 4 h (**Fig. 6E**), in accordance with the results obtained with CD8^+^ TILs from untreated and propranolol-treated mice (**Fig. 2C**).

**Figure 6.**
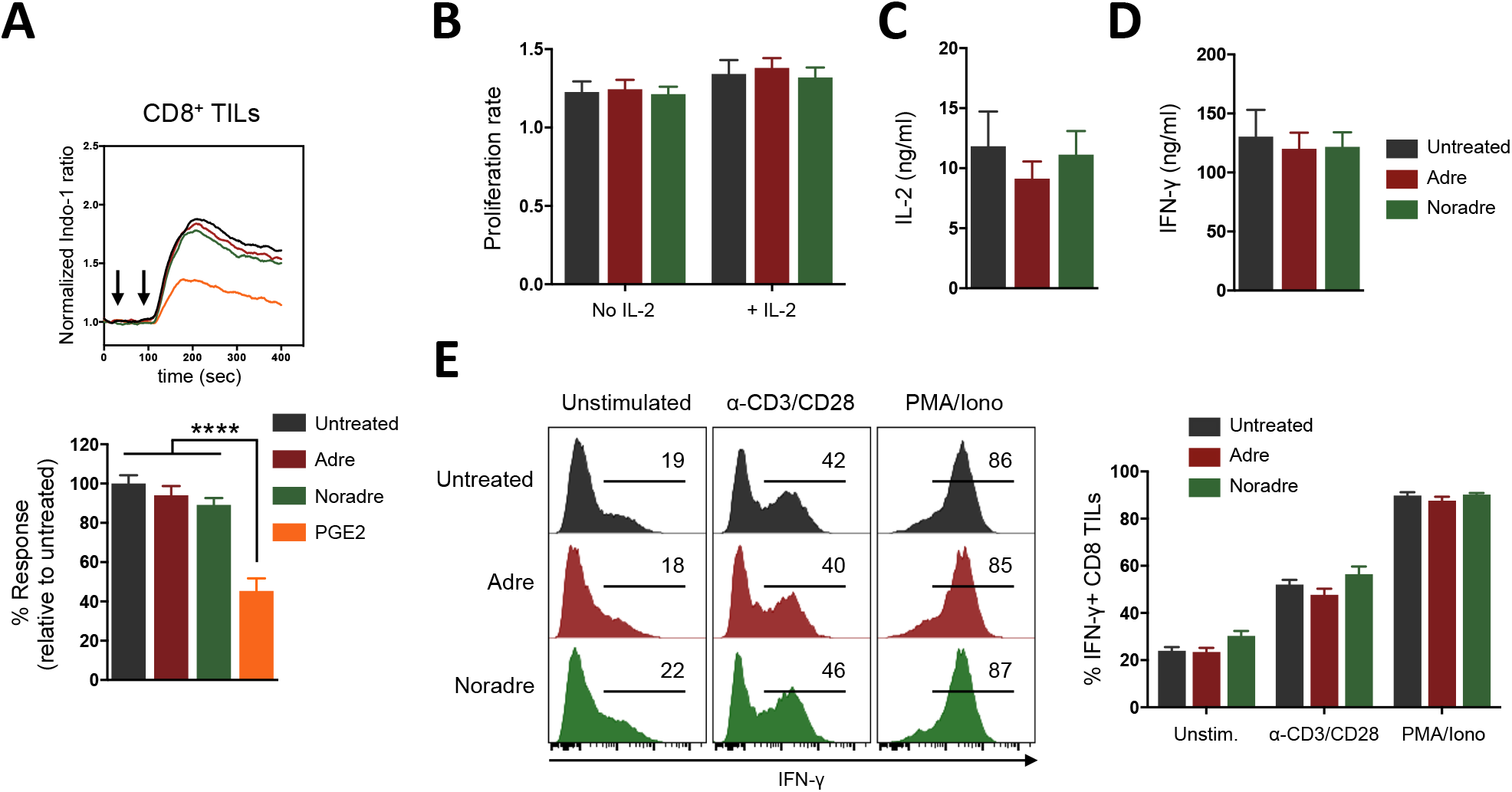
CD8^+^ TILs are not sensitive to adrenergic signals. **A**, Facs-sorted CD8^+^ TILs were loaded with Indo-1, pre-treated with either 10 μM adrenaline, noradrenaline or PGE_2_ during 10 min. iCa^2+^ mobilization was assessed by flow cytometry before and after stimulation with hamster anti-CD3ε Abs (first arrow, above) cross-linked with anti-hamster IgG (second arrow) and is indicated as % response in treated cells relative to untreated cells, with representative normalized Indo-1 ratio histograms. Pool from three independent experiments (*n =* 5). **B**, Proliferation rate of CD8^+^ TILs cultured in the presence of 10 μM adrenaline or noradrenaline with or without IL-2 (20 U/ml) during 24 h, represented as ratio of absolute number at day 0 / absolute number at day 1. Pool from two independent experiments (*n =* 9). **C** and **D**, IL-2 **(C)** and IFN-γ **(D)** measured in the supernatants 24 h after stimulation of CD8^+^ TILs cultured either alone or with IL-2 (20 Ul/ml) or restimulated with anti-CD3/CD28 Abs in the presence of 10 μM adrenaline or noradrenaline. Pool from two independent experiments (*n* = 5). E, Percentage of IFN-γ^+^ CD8^+^ TILs cells following *in vitro* restimulation with PMA/ionomycin or anti-CD3/CD28 Abs in the presence of 10 μM adrenaline or noradrenaline during 4 h, with representative flow cytometry histograms of intracellular IFN-γ staining in CD8^+^ T cells. Representative of two independent experiments *[n =* 5). Statistical analysis by one-way ANOVA test: ****, *p* <0.0001. Mean ± SEM.

### Differential sensitivity of naive CD8^+^ T cells and CD8^+^ TILs to β-AR signaling is related to their activation status and their β_2_-AR expression level

We wondered whether the differential sensitivity of naïve CD8^+^ T cells and CD8^+^ TILs to AR agonists could be related to their activation status. To test this hypothesis, we generated activated CD8^+^ T cells *in vitro* by stimulating them with anti-CD3/CD28 Abs during 3 days followed by a 4-day culture with IL-2. We then evaluated their sensitivity to adrenaline and noradrenaline using the same read-outs used with CD8^+^ TILs. As shown in **Fig 7A-D**, agonists had no effects on anti-CD3-induced calcium responses (**Fig. 7A**), proliferation rates (**Fig. 7B**) and IL-2/IFN-γ productions (**Fig. 7C, 7D**) by activated CD8^+^ T cells, similarly to what we had observed with CD8^+^ TILs. These results strongly suggest that activation of CD8^+^ T cells renders them insensitive to AR signaling, and that the refractory state observed with CD8^+^ TILs is likely a consequence of their activation status. In order to explain the difference between naive CD8^+^ T cells and CD8^+^ TILs/activated CD8^+^ T cells to AR signaling, we finally compared their β-AR expression at the mRNA levels. As for naive CD8^+^ T cells, CD8^+^ TILs and activated CD8^+^ T cells predominantly express β_2_-ARs (**Supplementary Fig. S7A, S7B**). However, the expression was much lower in CD8^+^ TILs and in activated CD8^+^ T cells, as compared to naive CD8^+^ T cells (**Fig. 7E**).

**Figure 7.**
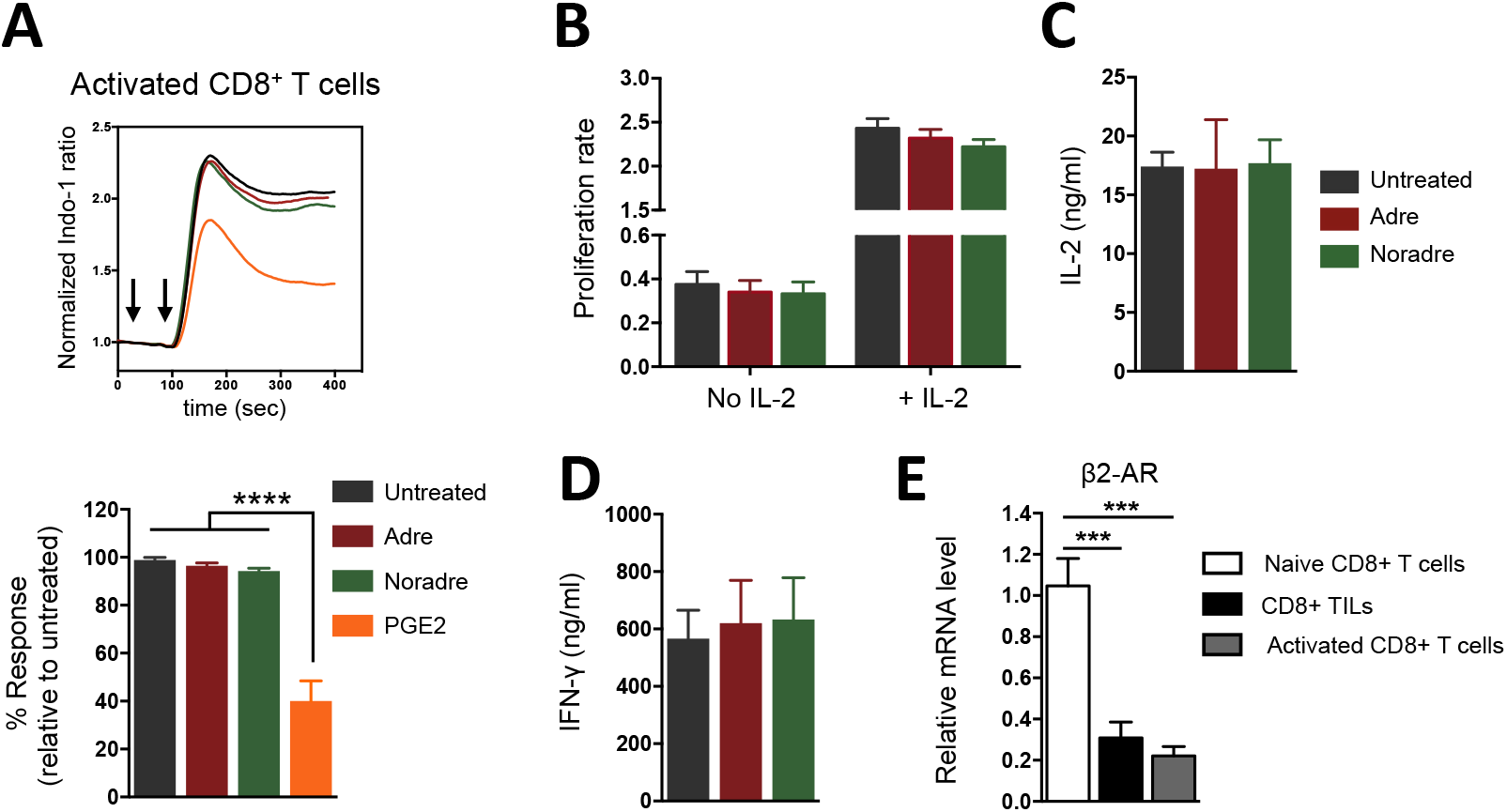
Differential sensitivity of naive CD8^+^ T cells and CD8^+^ TILs is related to their activation status and their β_2_-AR expression level. Activated CD8^+^T cells were obtained by stimulation with anti-CD3/CD28 Abs during 3 days following by 4 days cultured with IL-2. **A**, Activated CD8^+^ T cells were loaded with Indo-1, pre-treated with either 10 μM adrenaline, noradrenaline or PGE_2_ during 10 min. iCa^2+^ mobilization was assessed by flow cytometry before and after stimulation with hamster anti-CD3ε Abs (first arrow, above) cross-linked with anti-hamster IgG (second arrow) and is indicated as % response in treated cells relative to untreated cells, with representative normalized Indo-1 ratio histograms. Pool from four independent experiments (*n =* 4). **B**, Proliferation rate of activated CD8^+^ T cells cultured with or without IL-2 (20 UI/ml) in the presence of 10 μM adrenaline or noradrenaline during 24 h, represented as ratio of absolute number at day 0 / absolute number at day 1. Pool from two independent experiments (*n =* 9). **C** and **D**, IL-2 **(C)** and IFN-γ **(D)** measured in the supernatants 24 h after stimulation of activated CD8^+^ T cells with anti-CD3/CD28 Abs in the presence of 10 μM adrenaline or noradrenaline. Pool from two independent experiments (*n* = 4). (E) mRNA expression of β_2_-ARs in sorted-naive CD8^+^ T cells, CD8^+^ TILs and activated CD8^+^ T cells was evaluated by qPCR, (*n* = 5 to 6 per group). Statistical analysis by one-way ANOVA test: ***, *P* <0.001, ****, *p* <0.0001. Mean ± SEM.

Altogether these results demonstrate that the differential sensitivity of naive CD8^+^ T cells and CD8^+^ TILs to AR signaling is related to their activation status, most likely due to the down-regulation of the β_2_-AR expression driven by the activation.

## Discussion

In this study, we have shown that, in a therapeutic setting of vaccine-based immunotherapy designed for suboptimal efficacy, which mimics the suboptimal efficacy of cancer treatments for humans, a blockade of β-AR signaling by propranolol could markedly enhance the anti-tumor efficacy of the vaccine. This effect was due to a large extent to an increased tumor CD8^+^ T cell infiltrate. A crucial part of the beneficial effect of propranolol was an improved efficacy of the vaccine at the priming phase against tumor antigen in the TDLN, likely due to its direct action on naive CD8^+^ T cells. We have also examined if the effector phase of the anti-cancer response by CD8^+^ TILs within the tumor was affected. To our surprise, we did not observe any obvious influence of propranolol on the reactivity of CD8^+^ TILs, a result further reinforced by *ex-vivo* experiments showing that, contrary to naive CD8^+^ T cells, CD8^+^ TILs were insensitive to AR signaling. Finally, we have shown that this lack of response of CD8^+^ TILs to β-AR signaling is likely related to their activation phenotype, characterized by a down-regulation of β_2_-AR expression.

Over the last decade, several studies have revealed that β-AR signaling could facilitate tumor growth and cancer progression (14,15,20,33). Many of them were performed in immunodeficient mice (xenograft tumor models), focusing on a direct action of β-AR signaling on tumor cells or on pro-tumorigenic processes such as angiogenesis. However, in such experimental settings, the impact of β-AR signaling on the associated anti-tumor immune response has been unavoidably underestimated, as it is now clear that the autonomic nervous system may have an important role in regulating immune responses (34). In most of the studies seeking to understand the role of the sympathetic system, adrenergic signals were generated by applying a specific chronic psychological stress stimulus (*e.g.* restraint stress, social defeat stress, social isolation). Our choice has been to work in the least artificial situation, *i.e.* just under baseline housing stress. Indeed, the housing conditions of our mice, such as sub-thermoneutral housing (20-26°C), have been described as a physiologic model of mild stress generating chronic adrenergic signaling (35). Moreover, in some studies, propranolol treatment was given prior to tumor implantation and it has been demonstrated that the ability of immune responses to control a tumor was different whether treatment was given prior or after tumor implantation (26). We chose to treat mice when their tumor reached a palpable size, considering such a therapeutic approach as most relevant clinically. Finally, the TC1 model used in our study presented the advantage that propranolol had no effect on tumor growth in the absence of vaccination, a condition that is quite helpful for describing precisely the effects of propranolol blockade on a vaccine-induced anti-tumor immune response.

We found that the major effect of β-AR signaling blockade by propranolol treatment occurred at the CD8^+^ T-cell response level. Our findings are consistent with recent studies showing that adrenergic signals impaired the anti-tumor CD8^+^ T-cell response, a phenomenon that may be reversed by β-blocker treatment (26–28,36). Interestingly, the inhibition of the CD8^+^ T-cell response by adrenergic signals is not limited to the tumor setting and has also been shown in the context of infection, such as influenza (37). Obviously, these findings do not imply that β-AR signaling cannot affect other immune cell populations, in particular myeloid cells. Indeed, β-AR signaling has been shown to promote tumor growth or metastasis through M2 polarization of macrophages (19,20) and infiltration of MDSCs (21,22). However, we did not observe in our tumor model differences in the frequency of myeloid cells, in accordance with the fact that chemokine production was also not affected by propranolol treatment. Moreover, we did not observe any obvious effect of propranolol on the activation state of myeloid cells, reflected by their MHC-II expression level (data not shown). Nevertheless, future work would deserve to be done to specifically explore in more details the potential effects of β-AR signaling on the suppressive functionality of MDSCs and TAMs in this model.

Our study provides an additional insight into the beneficial effect of β-AR blockade by showing that it can potentiate the priming of naive CD8^+^ T cells that takes place in the TDLN. We found a strong inhibitory effect of β-AR signaling on the proliferation of naive CD8^+^ T cells. This result suggests a direct inhibitory effect on CD8^+^ T cells during their priming phase in the TDLN, which could explain the higher frequency of specific CD8^+^ T cells in TDLN and the tumor in vaccinated mice treated with propranolol. However, we do not exclude the possibility of additional mechanisms. Indeed, previous studies have showed that adrenergic signals reduced the antigen presentation, maturation, cytokine production and migration of DCs (38–42) that could indirectly affect the T-cell priming. In a recent paper, Sommershof and colleagues have shown that chronic stress suppressed anti-tumor CD8^+^ T-cell response following anti-cancer vaccination by impairing DC maturation, migration and subsequent CD8^+^ T-cell priming (28). However, in our tumor vaccine model, we showed that adrenergic signals did not affect the maturation and the antigen presentation capacities of vaccine-stimulated DCs. This is in accordance with recent studies showing that DC maturation was not affected by β_2_-AR signaling (42) and that its inhibitory effect on CD8^+^ T-cell response induced by immune modulating Abs occurred independently of changes in DC functions (27). Another possibility is an effect of propranolol on lymphocytes egress from LNs, as this process has been shown to be controlled by β_2_-ARs (43). If this point was relevant in our model, the increased frequency of CD8^+^ TILs we observed would be the result of an enhanced egress from the TDLN induced by β_2_-AR blockade. In this case, we would expect to have a lower frequency of specific CD8^+^ T cells in the TDLN from propranolol-treated vaccinated mice. On the contrary, we clearly observed a higher frequency, suggesting that a decisive part of the potentiating effect of propranolol operates through the priming phase of T-cell activation, after TCR engagement.

A major and surprising observation of our study is the differential sensitivity of naive versus TILs/activated CD8^+^ T cells to AR agonists. We found that, contrary to naive CD8^+^ T cells, CD8^+^ TILs and activated CD8^+^ T cells were insensitive to adrenaline/noradrenaline treatment, and this was associated with a down-regulation of the β_2_-AR mRNA expression. Our data are consistent with previous studies reporting differential β_2_-AR expression in T-cell subsets, with implications on their functions. Thus, Nakai and colleagues have shown that murine naive CD4^+^ T cells expressed more β_2_-ARs than effector/memory CD4^+^ T cells (43). On the contrary, other studies have observed that human memory CD8^+^ T cells expressed more β_2_-ARs than naive CD8^+^ T cells (44,45). Tregs were also shown to express lower level of β_2_-ARs than naive CD4^+^ T cells (46). Moreover, it has been shown that CD4^+^ T-cell polarization towards a Th2 profile was accompanied by a loss of β_2_-AR expression (47,48), as a result of epigenetic regulation (49). Could similar epigenetic regulation occur during activation of naive CD8^+^ T cells leading to the down-regulation of β_2_-ARs in CD8^+^ TILs and activated CD8^+^ T cells is therefore an important question that remains to be addressed.

Several questions and potential future studies emerge from this work. What are the exact molecular mechanisms underlying the inhibition of T-cell activation by β_2_-AR signaling? To our knowledge, we provide here the first demonstration that a CD3-induced calcium response is inhibited by adrenaline or noradrenaline treatment in naive CD8^+^ T cells. This shows that the inhibition takes place at a very early stage of the TCR signaling pathway. Engagement of the β_2_-AR is known to activate adenylate cyclase, catalyzing the formation of cAMP, which activates PKA. cAMP is a well known inhibitor of T-cell activation (50). PKA-mediated activation of Csk, a major regulator of Src kinases activity, was demonstrated as one mechanism for cAMP-dependent inhibition of lymphocyte activation (51). However, other PKA-independent signaling pathways, such as the guanine nucleotide exchange protein activated by adenylyl cyclase (EPAC) and β-arrestin can also be activated by β_2_-AR engagement. Indeed, in DCs, Takerana and colleague have recently showed that down-regulation of IL-12p70 mediated by β_2_-AR activation was related to the inhibition of NF-kB activation and AP-1 signaling, at least partially dependent on β-arrestin (42). Thus, the exact contribution of these different pathways in the T-cell inhibition induced by β_2_-AR engagement remains to be investigated.

What are the clinical implications of our findings? These last years, immunotherapies have emerged as a great promise in cancer treatment. However, many tumor types are not responding yet and, even for sensitive tumors, the mechanisms limiting cancer immunotherapies deserve to be elucidated in more details. Therefore, finding new strategies aiming at improving the efficacy of cancer immunotherapies is a crucial objective. One of them is to identify immunosuppressive factors of the TME interfering with these new treatments in order to reverse their negative impact. In our study, we demonstrate that an induced anti-tumor CD8^+^ T-cell response can be greatly improved by β-blockers removing the detrimental effects of endogenous catecholamines. These findings are in accordance with studies showing that β_2_-AR signaling decreases the efficacy of several immunotherapeutic approaches, including therapies using immune checkpoint inhibitors (26–28). They are also in line with retrospective studies suggesting that cancer patients taking β-blockers have better outcomes (11–13). Importantly, we also introduce a new major notion by demonstrating that β-AR blockade may act as an adjuvant during the priming of the anti-tumor CD8^+^ T-cell response. These results strengthen the rationale for using inexpensive and well-tolerated β-blockers in clinical oncology for potentiating some immunotherapies. In particular, those triggering an anti-tumor T-cell response, either by a specific vaccine, or after a treatment inducing an immunogenic tumor cell death leading, in the TDLNs, to the presentation of tumor antigens and the priming of naive CD8^+^ T cells. Finally, these results support the idea that psychological stress, which is a powerful provider of adrenergic signals, must be taken into account and managed when treating cancer patients.

## Supporting information

## Acknowledgements

We wish to thank the imaging (IMAG’IC), flow cytometry (CYBIO) and genomic (GENOM’IC) facilities of the Cochin Institute for their helpful advice, people from “Dynamic of T cell interactions” team for their helpful discussions. We are grateful to Agnes Le Bon for providing IFN-α and Eric Tartour and Judger Johannes for providing Shiga vaccines. This study was supported by grant from the French Ligue Nationale contre le Cancer (VF). Clara Daher was supported by a PhD fellowship from the French Ministry of National Education, Research, and Technology.

## Authors’ Contributions

Conception and design: C. Daher, V. Feuillet

Development of methodology: C. Daher, L. Vimeux, N. Bercovici, V. Feuillet

Acquisition of data (provided animals, acquired and managed patients, provided facilities, etc.): C. Daher, L. Vimeux, R. Stoeva, E. Peranzoni, V. Feuillet

Analysis and interpretation of data (e.g., statistical analysis, biostatistics, computational analysis): C. Daher, L. Vimeux, R. Stoeva, V. Feuillet

Writing, review, and/or revision of the manuscript: C. Daher, L. Vimeux, E. Peranzoni, G. Bismuth, E. Donnadieu, N. Bercovici, A. Trautmann, V. Feuillet

Administrative, technical, or material support (i.e., reporting or organizing data, constructing databases): C. Daher, L. Vimeux, E. Donnadieu, V. Feuillet

Study supervision: V. Feuillet

## Abbreviations

STxB: nontoxic B subunit of Shiga toxin
TIL: tumor-infiltrating lymphocyte
AR: adrenergic receptor
DC: dendritic cell
TDLN: tumor-draining lymph node
TAM: tumor-associated macrophage
PMN: polymorphonuclear cell
NK cell: natural killer cell
Treg: regulatory CD4^+^ T cell
MDSC: myeloid derived suppressive cell
TME: tumor microenvironment
PGE2: prostaglandin E2
BMDC: bone marrow derived dendritic cell
Abs: antibodies
PKA: protein kinase A

