## Supplementary material for "Blockade of β-adrenergic receptor signaling improves cancer vaccine efficacy through its effect on naive CD8^+^ T-cell priming"

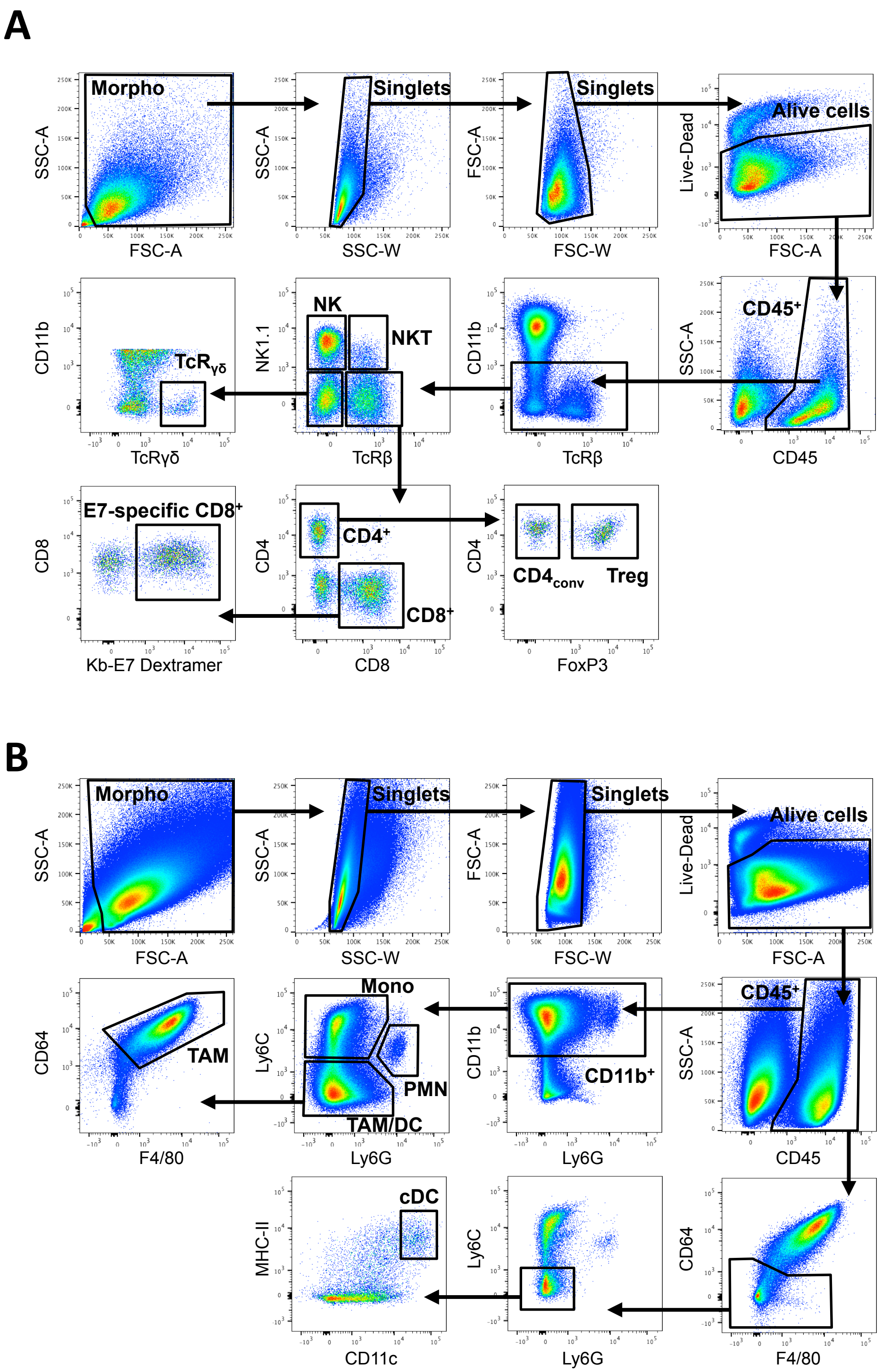


**Figure S1. Gating strategy for flow cytometry analysis of the immune infiltrate in the tumor**. Cells were gated based on their forward (size) vs side (granulosity) scatter. After two steps of cell-doublet exclusion, dead cells were gated out and cells were gated based on their CD45 expression, sequentially. **A,** Gating strategy for flow cytometry analysis of the lymphoid compartment infiltrate in the tumor. $CD45^{+}$-lymphoid cells were defined as followed: NK cells ($NK1.1^{+}TcR\beta^{neg}$), NKT cells ($NK1.1^{+}TcR\beta^{+}$), $T\gamma\delta^{+}$ cells ($NK1.1^{neg}TcR\beta^{neg}TcR\gamma\delta^{+}$), regulatory $CD4^{+}$ T cells (Treg; $NK1.1^{neg}TcR\beta^{+}CD4^{+}FoxP3^{+}$), conventional $CD4^{+}$ T cells (CD4conv; $NK1.1^{neg}TcR\beta^{+}CD4^{+}FoxP3^{neg}$), $CD8^{+}$ T cells ($NK1.1^{neg}TcR\beta^{+}CD8^{+}$), E7-specific $CD8^{+}$ T cells ($NK1.1^{neg}TcR\beta^{+}CD8^{+}$Kb-E7-dextramer$^{+}$). **B,** Gating strategy for flow cytometry analysis of the myeloid compartment infiltrate in the tumor. $CD45^{+}$-myeloid cells were defined as followed: monocytes (Mono; $CD11b^{+}Ly6G^{neg}Ly6C^{+}$), Polymorphonuclear Neutrophils (PMN; $CD11b^{+}Ly6G^{+}Ly6C^{int}$), Tumor-Associated Macrophages (TAM; $CD11b^{+}Ly6G^{neg}Ly6C^{neg}CD64^{+}F4/80^{+}$), conventional Dendritic Cells (cDC; $CD64^{neg}F4/80^{neg}Ly6G^{neg}Ly6C^{neg}CD11c^{+}MHC\text{-}II^{+}$).
