## Supplementary material for "Blockade of β-adrenergic receptor signaling improves cancer vaccine efficacy through its effect on naive CD8^+^ T-cell priming"

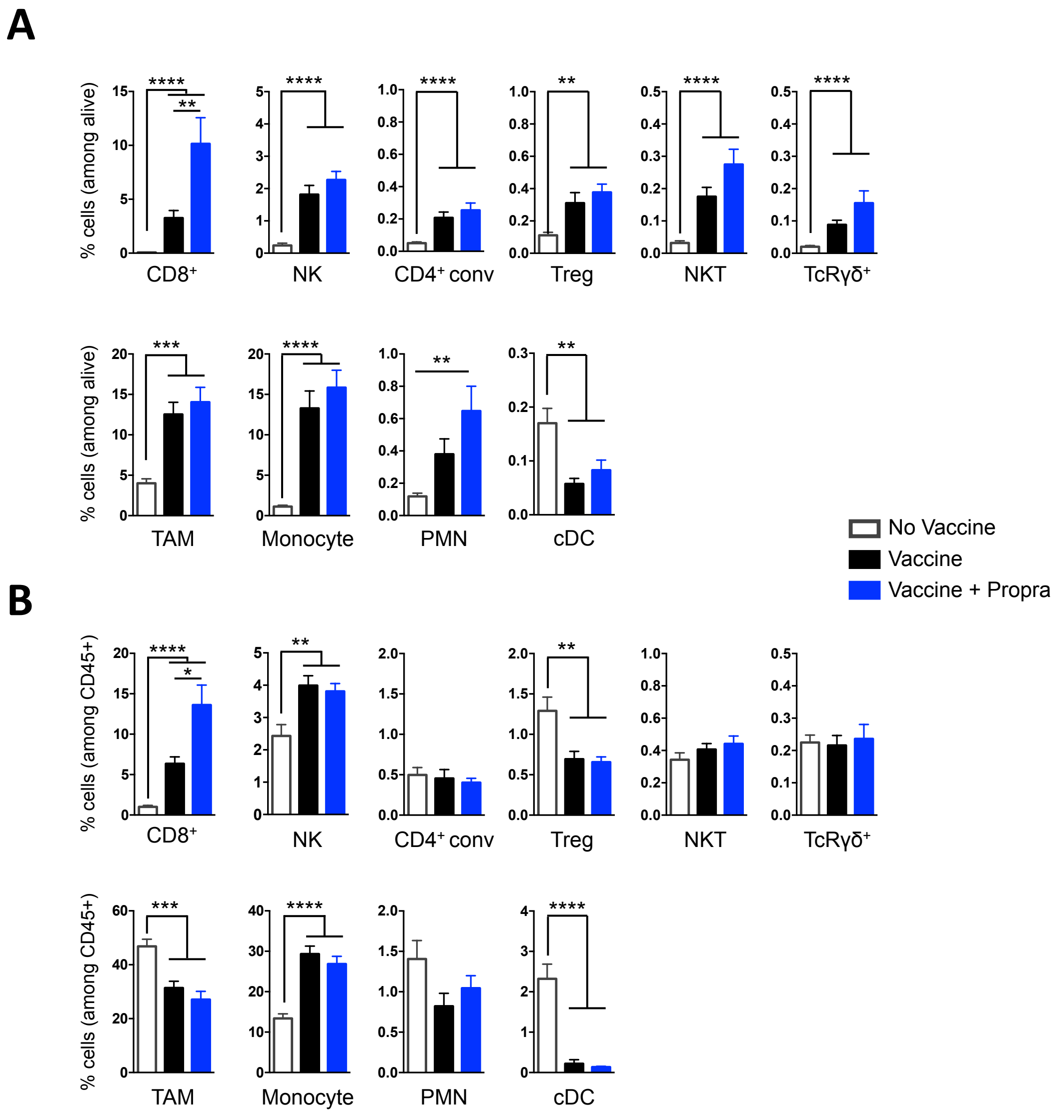


**Figure S2. Analysis of the immune infiltrate composition** C57BL/6J mice were s.c. injected with E7 expressing-TC1 tumor cells and vaccinated as mentioned in Figure 1. Day 10 post-vaccination, frequencies of immune cells (defined in Supplementary Figure 1) among live cells (**A**) and among CD45+-cells (**B**) from tumor single-cell suspensions evaluated by flow cytometry. Pool from six independent experiments (*n* = 16 to 18 per group). Statistical analysis by Mann-Whitney test: *, *P* <0.05, **, *P* <0.01, ***, *P* <0.001, ****, *P* <0.0001. Mean ± SEM.
