## Supplementary material for "Blockade of β-adrenergic receptor signaling improves cancer vaccine efficacy through its effect on naive CD8^+^ T-cell priming"

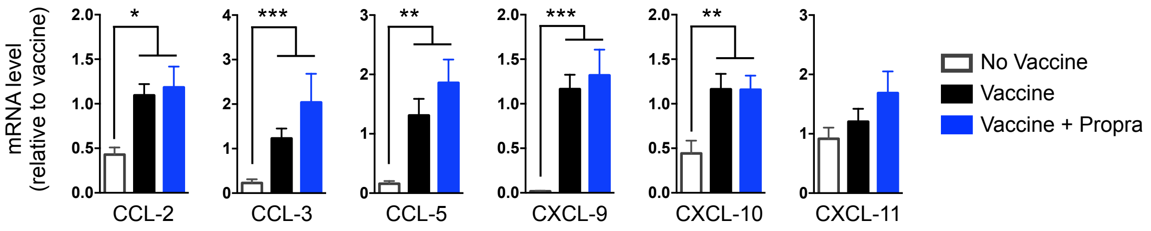


**Figure S3. Chemokine mRNA transcripts in the tumor.** C57BL/6J mice were s.c. injected with E7 expressing-TC1 tumor cells and vaccinated as mentioned in Figure 1. Day 10 post-vaccination, mRNA expression of indicated chemokines in whole tumor was evaluated by qPCR. Pool from four independent experiments (*n* = 7 to 14 per group). Statistical analysis by Mann-Whitney test: *, *P* <0.05, **, *P* <0.01, ***, *P* <0.001. Mean ± SEM.
