## Supplementary material for "Blockade of β-adrenergic receptor signaling improves cancer vaccine efficacy through its effect on naive CD8^+^ T-cell priming"

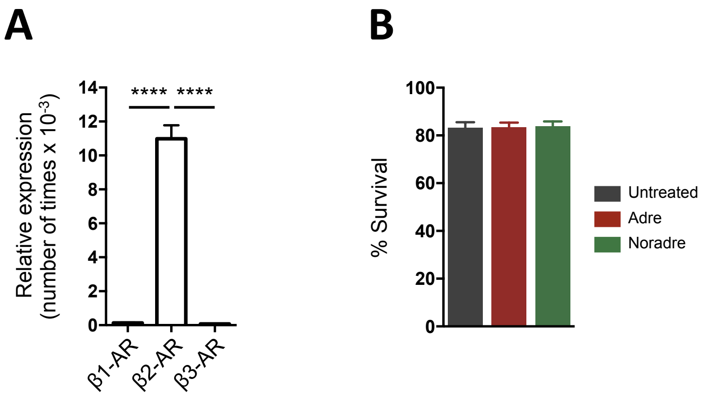


**Figure S4. mRNA expression of β1-, β2-, β3-ARs in naive CD8+ T cells and effect of adrenaline/noradrenaline treatment on naive CD8+ T-cell viability. A,** mRNA expression of β1-, β2-, β3-ARs was evaluated by qPCR. Bars show the number of times that target genes are expressed as compared to the housekeeping gene GADPH in Facs-sorted naive CD8+ T cells. Pool from 3 independent experiments (*n* = 6). **B,** Viability of naive CD8+ T cells cultured with IL-7 (10 ng/ml) in the presence of 10 µM adrenaline or noradrenaline for 72 h, evaluated with Live/dead staining by flow cytometry. Pool from three independent experiments, (*n* = 13). Statistical analysis by one-way ANOVA test: ****, $P < 0.0001$. Mean ± SEM.
