## Supplementary material for "Blockade of β-adrenergic receptor signaling improves cancer vaccine efficacy through its effect on naive CD8^+^ T-cell priming"

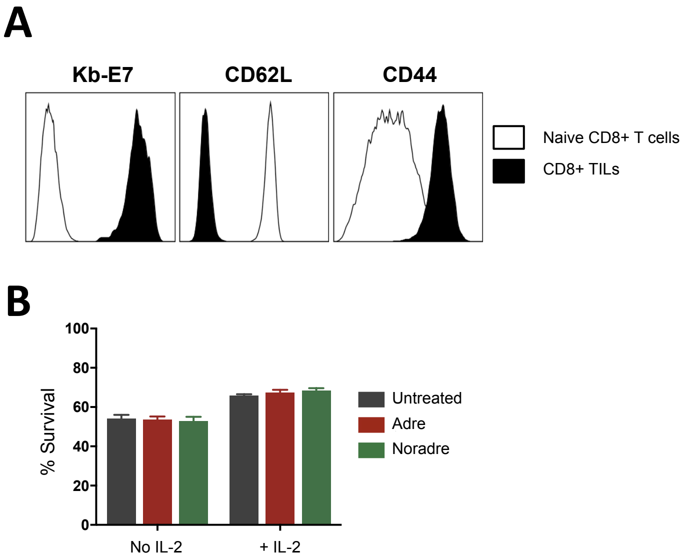


**Figure S5. Activation status of flow-cytometry sorted CD8⁺ TILs and effect of adrenaline/noradrenaline treatment on CD8⁺ TIL viability. A,** Expression of CD44, CD62L and Kb-E7 dextramer staining on Facs-sorted $CD8^+$ TILs from vaccinated tumors and naive $CD8^+$ T cells (as negative control) evaluated by flow cytometry. **B,** Viability of $CD8^+$ TILs cultured with or without IL-2 (20 U/ml) in the presence of 10 µM adrenaline or noradrenaline during 24 h, evaluated with Live/dead staining by flow cytometry. Pool from two independent experiments ($n$ = 12). Statistical analysis by one-way ANOVA test. Mean ± SEM.
