## Supplementary material for "Blockade of β-adrenergic receptor signaling improves cancer vaccine efficacy through its effect on naive CD8^+^ T-cell priming"

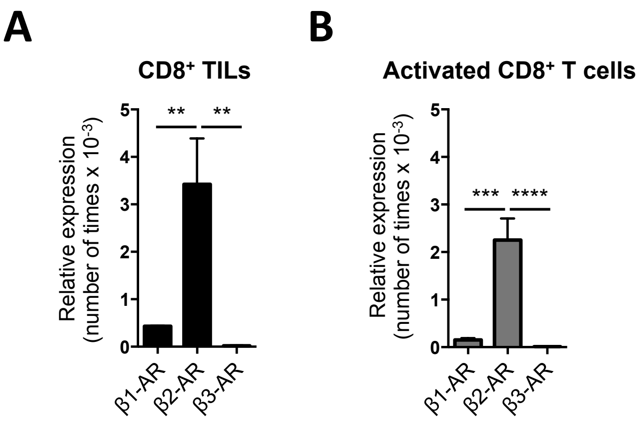


**Figure S6. mRNA expression of β1-, β2-, β3-ARs in CD8⁺ TILs and activated CD8⁺ T cells.** mRNA expression of β1-, β2-, β3-ARs was evaluated by qPCR. Bars show the number of times that target genes are expressed as compared to the housekeeping gene GADPH in CD8⁺ TILs (**A**) and activated CD8⁺ T cells (**B**). Pool from 3 independent experiments, (*n* = 5 to 6 per group). Statistical analysis by one-way ANOVA test: **, *P* <0.01, ***, *P* <0.001, ****, *P* <0.0001. Mean ± SEM.
