## Supplementary material for "Blockade of β-adrenergic receptor signaling improves cancer vaccine efficacy through its effect on naive CD8^+^ T-cell priming"

| Genes | Forward primers | Reverse primers |
|---|---|---|
| **CCL-2** | CATCCACGTGTTGGCTCA | GATCATCTTGCTGGTGAATGAGT |
| **CCL-3** | TGCCCTTGCTGTTCTTCTCT | GATGAATTGGCGTGGAATCT |
| **CCL-5** | AGCAGCAAGTGCTCCAATCT | ATTTCTTGGGTTTGCTGTGC |
| **CXCL-9** | TCCTTTTGGGCATCATCTTCC | TTTGTAGTGGATCGTGCCTCG |
| **CXCL-10** | TCCTTGTCCTCCCTAGCTCA | ATAACCCCTTGGGAAGATGG |
| **CXCL-11** | GGCTTCCTTATGTTCAAACAGGG | GCCGTTACTCGGGTAAATTACA |
| **β1-AR** | GCTGATCTGGTCATGGGATT | AAGTCCAGAGCTCGCAGAAG |
| **β2-AR** | GGGAACGACAGCGACTTCTT | GCCAGGACGATAACCGACAT |
| **β3-AR** | CGAAGAGCATCA CAAGGAGGG | CGAAACTGGTTGCGGAACTGTGT |
| **GAPDH** | TGTGTCCGTCGTGGATCTGA | TTGCTGTTGAAGTCGCAG |

**Table S1. Primer sequences for RT-qPCR.**
